# Enabling the hypothesis-driven prioritization of ligand candidates in big databases: Screenlamp and its application to GPCR inhibitor discovery for invasive species control

**DOI:** 10.1101/249151

**Authors:** Sebastian Raschka, Anne M. Scott, Nan Liu, Santosh Gunturu, Mar Huertas, Weiming Li, Leslie A. Kuhn

**Affiliations:** Department of Biochemistry and Molecular Biology, Michigan State University, East Lansing, MI 48824, USA; Department of Fisheries and Wildlife, Michigan State University, East Lansing, MI 48824, USA; Department of Chemistry, Michigan State University, East Lansing, MI 48824, USA; Department of Computer Science and Engineering, Michigan State University, East Lansing, MI 48824, USA; Current address: Department of Biology, Texas State University, San Marcos, TX 78666, USA

**Keywords:** virtual screening, structure based drug discovery, G protein coupled receptor, chemoinformatics, computer-aided molecular design, structure-activity relationships

## Abstract

While the advantage of screening vast databases of molecules to cover greater molecular diversity is often mentioned, in reality, only a few studies have been published demonstrating inhibitor discovery by screening more than a million compounds for features that mimic a known three-dimensional ligand. Two factors contribute: the general difficulty of discovering potent inhibitors, and the lack of free, user-friendly software to incorporate project-specific knowledge and user hypotheses into 3D ligand-based screening. The Screenlamp modular toolkit presented here was developed with these needs in mind. We show Screenlamp’s ability to screen more than 12 million commercially available molecules and identify potent *in vivo* inhibitors of a G protein-coupled bile acid receptor within the first year of a discovery project. This pheromone receptor governs sea lamprey reproductive behavior, and to our knowledge, this project is the first to establish the efficacy of computational screening in discovering lead compounds for aquatic invasive species control. Significant enhancement in activity came from selecting compounds based on one of the hypotheses: that matching two distal oxygen groups in the three-dimensional structure of the pheromone is crucial for activity. Six of the 15 most active compounds met these criteria. A second hypothesis – that presence of an alkyl sulfate side chain results in high activity – identified another 6 compounds in the top 10, demonstrating the significant benefits of hypothesis-driven screening.

## Introduction

### Virtual screening for inhibitor discovery

Virtual screening in biology employs computational algorithms to filter or compare the structures or chemical features of thousands to millions of compounds to uncover those similar to a known bioactive reference molecule, or to identify a subset of compounds with significant complementarity to a receptor-binding site. One goal of screening is to provide a way for the biologist to encode and test a model of the features he hypothesizes to be important in small molecules that inhibit or activate the chosen receptor. Computational screening also leverages, on a scale not accessible to most experimental screening groups, the ability to test many or all of the millions of small drug-like molecules that are now available commercially. The much more expensive and labor-intensive lab assays can then be performed on a subset of screening-identified molecules with a significantly enhanced probability of identifying molecules with the desired bioactivity. Screening typically has a good success rate, with 5-35% of the resulting prioritized and tested molecules proving to be moderate to high affinity inhibitors, whereas high-throughput experimental screening involving no computational hypothesis-based prioritization typically has a success rate of 0.01% (1 active in every 10,000 molecules tested) [1, 2]. Here, we present a virtual screening software framework, Screenlamp, used to identify inhibitors of a known modulator of G protein-coupled receptor signaling, by combining unbiased 3-dimensional shape and chemical similarity alignment with the ability to test investigator-formulated hypotheses about chemical groups that lead to inhibitor or activator activity. This software facilitates the use of one or more of the three commonly used approaches: (1) comparison of molecules in a database for similarity with a known bioactive molecule, (2) definition of the spatial relationships between chemical groups in a small molecule that allow it to activate or inhibit a biological receptor (known as pharmacophore identification and matching), and (3) docking of small molecules into a receptor structure to identify the subset of molecules that interact most favorably with the receptor.

In receptor structure-based approaches, small molecules are docked into the three-dimensional (3D) structure of an enzyme or receptor binding site to select a set of molecules for experimental testing as activators or inhibitors of the protein. The prioritization of candidates is typically based on ranking the molecules by their predicted binding affinities [3]. However, applications of structure-based screening are limited by the availability of accurate three-dimensional (3D) structures of the target protein. Moreover, a consequence of the large number of geometrically feasible solutions when both molecules are considered flexible is that thorough sampling of such docking poses is computationally impractical, even for state-of-the-art computing clusters. As a result, most currently used docking solutions treat the ligand candidate as flexible and the protein as only partly flexible via limited side-chain sampling [4]. Even under these partially-flexible protein assumptions, ligand docking is very computationally expensive. It is not feasible for most academic research groups to dock millions of small, flexible molecules, which requires the use of computing clusters or commercial cloud services [5]. An equally significant problem is that prediction of ΔG_binding_ of protein-ligand complexes has remained prone to errors typically on the order of several kcal/mol (a substantial percentage of the total ΔG_binding_), causing the ranking of compounds to be approximate at best [6]. This problem is likely to remain difficult and improve incrementally rather than rapidly, due to the difficulty of measuring conformational energies, entropy changes, electrostatics, and solvent contributions to ligand-receptor binding [7]. The most accurate approaches are only feasible for assessing a small number of compounds.

Ligand-based screening, in which database compounds are compared to a known active compound (rather than docked to the protein target) to discover mimics, is frequently employed by pharmaceutical companies due to the success rate and the unavailability of 3D protein structures for many targets of interest. Generally, ligand-based virtual screening is computationally more efficient than structure-based approaches [8]. An additional advantage is that errors in modeling protein and solvent flexibility do not come into play in 3D ligand similarity-based scoring, which is based solely on the extent to which a candidate matches the known ligand in volume and charge or atom-type distribution. Ligand-based screening can outperform structure-based approaches in the speed and the enrichment of active molecules [9–11]. Furthermore, when performed with a single known active compound for comparison, 3D ligand-based screening is capable of identifying molecular mimics with diverse structural scaffolds and chemotypes. This desirable feature, known as lead or scaffold hopping [12, 13], is important since a significant percentage of inhibitory compounds may undergo attrition during the pharmacological and clinical development process due to not meeting criteria for *in vivo* absorption, distribution, metabolism, excretion, and toxicity.

The Screenlamp project started when we sought to fill a gap, by developing freely available, effective software to enable typical academic biochemical research groups, rather than just computational chemistry experts, to test their hypotheses about the importance of specific functional groups or pharmacophores (3D spatial relationships between functional groups) that lead to high ligand activity, when performing a broad search for compounds or scaffolds with significant similarity to a known ligand. In a random database, the probability of finding one or more good lead molecules with substantial affinity for the protein target via close mimicry of a known ligand increases with the number of molecules screened. Thus, our second goal was to make the organization and screening of very large databases of millions of commercially available compounds accessible to a typical research lab, rather than being restricted to researchers with parallel computing expertise. Some tools exist to aid users in ligand-based screening, but they are limited by the level of molecular detail they support, the flexibility of use, and cost. The SwissSimilarity webserver was recently launched to support ligand-based virtual screening [14]. While this service includes 10.6 million drug-like molecules from ZINC, its screening is based on non-superpositional methods that do not consider the 3D volumes or spatial arrangement of functional groups. Phase is a commercial tool developed by Schrödinger, which allows users to perform 3D ligand-based screening based on abstract hydrogen-bond acceptor and donor, hydrophobic, aromatic, and charged pharmacophore points, which the software derives from known actives [15]. Aside from the barrier of substantial licensing costs, its integration as part of the Schrödinger graphical user interface, including assignment of ligand protonation states and conformers and the use of a proprietary scoring function and eMolecules database, limits its flexibility. We have found partial charge and protonation state assignment, quality of 3D conformer sampling, flexible identification of pharmacophores and querying based on functional group relationships, and 3D overlays and similarity scoring to be variable in quality between existing software packages while being essential to screening success. The ability to choose the modules that work best for a project and provide a freely available, flexible workflow for 3D ligand-based discovery are supported by Screenlamp.

The fact that 3D ligand-based screening of millions of compounds still presents a substantial technical challenge to most users is underscored by only a few inhibitor-discovery publications appearing in the literature for this approach over the past 13 years [16–25], in comparison with dozens of publications for screening by docking of similar-sized databases. With Screenlamp, volumetric and partial charge-based alignment of fully flexible molecules and analysis of 3D chemical group matches can be performed on millions of commercially available molecules, such as the ZINC drug-like database [26] (http://zinc.docking.org), within a day on a typical desktop computer. The Screenlamp toolkit provides a set of command line virtual screening utilities that can be used individually to perform specific filtering tasks or in an automated fashion as a computational pipeline (with examples and templates provided in the distribution package available at https://psa-lab.github.io/screenlamp/) on desktop computers or computing clusters. Here we demonstrate its successful application to a challenging problem: discovery of both steroidal and non-steroidal inhibitors with IC50 values under 1 μM for an olfactory GPCR activated by a bile acid pheromone [27]. Because the molecular weight of the pheromone is at the upper limit for drug-like compounds, the discovery of active compounds benefited from Screenlamp’s ability to search expanded sets of molecules from ZINC and the Chemical Abstracts Service Registry (https://www.cas.org/content/chemical-substances).

### Pioneering aquatic invasive species control and GPCR inhibitor discovery with Screenlamp

This pheromone inhibitor discovery project presents a novel, behaviorally selective approach to aquatic invasive species control, which in the past has involved *in vivo* testing of thousands of pesticides. The sea lamprey is an invasive species that has had greatly deleterious impacts since the 1950s on both the native ecology and commercial fishery of the Laurentian Great Lakes. Ongoing efforts at reducing sea lamprey populations are labor-intensive and cost millions of dollars per year [28]. They include the use of in-stream barriers to prevent lamprey from reaching spawning areas [29] and the application of trifluoromethyl nitrophenol (TFM), a larval lampricide [30]. TFM has been successful, leading to a decrease of the sea lamprey population by over 90% between 1960 and 1970 [31]. However, the discovery of new sea lamprey control approaches remains a high priority for the binational Great Lakes Fishery Commission. Occasionally, TFM has shown off-target toxic effects to amphibians, trout, and most importantly, lake sturgeon, which the U.S. Fish and Wildlife Service lists as threatened or endangered in nineteen of the twenty states of its historic range (https://www.fws.gov/midwest/sturgeon/biology.htm; date accessed: Sept. 2, 2017) [32, 33]. A recent sea lamprey control approach involves baiting traps [34, 35] with the main component of the male sea lamprey mating pheromone 3kPZS (3-keto petromyzonol sulfate; 7α,12α,24-trihydroxy-5α-cholan-3-one-24-sulfate), which is an agonist for the sea lamprey odorant receptor 1 (SLOR1).

### G-protein coupled receptors and olfactory receptors

SLOR1 (UniProtKB ID: S4RTH2) and other pheromone and olfactory receptors in the sea lamprey are categorized as class A G protein-coupled receptors (GPCRs) based on sequence homology [36]. Class A or rhodopsin-like GPCRs form the largest of the five GPCR superfamilies [37]. GPCRs play an important role in human medicine, with about half of all human drugs targeting GPCRs and their signaling [38]. A well-known agonist of β1-adrenergic receptor, the closest human structural homolog of SLOR1, is epinephrine (also known as adrenaline). Antagonists of this receptor known as beta blockers are commonly used for controlling blood pressure and glaucoma. In humans, olfactory receptors comprise 388 out of our 779 GPCRs (http://gpcr.usc.edu), indicating their importance for responding to chemical cues in the environment, such as oxygen [39], smoke [40], scents released by rotten meat [41] or associated with nutrients [42], and pheromones [43]. Ligands for class A GPCRs are correspondingly diverse, including steroids, peptides, light-responsive chromophores, neurotransmitters, lipids, nucleotides, and chemokine proteins [44]. In insects, non-GPCR olfactory receptors [45] also play key roles in sensing and responding to repellants such as DEET, as well as in detecting pheromones that lead to mating and reproduction [46].

Sea lamprey mating is governed by sex pheromones released by spermiated mature males. Ovulated females are attracted by the 3kPZS secreted in spawning areas. We hypothesized that blocking the detection of 3kPZS by female sea lamprey would halt the reproductive cycle and reduce the sea lamprey population. The aim of our high-throughput screening was thus to identify sea lamprey-selective inhibitor mimics of 3kPZS that are environmentally benign. Screenlamp was developed and used to screen 12 million commercially available small organic molecules. Those with the most significant volumetric and atom-type similarity to 3kPZS were further prioritized within Screenlamp by filtering compounds according to a series of hypotheses about the importance of individual chemical groups for activity. *In vivo* olfactory assays of the selected 299 compounds were then performed, testing their ability to block 3kPZS olfactory responses in sea lamprey, and resulting in the discovery of several classes of inhibitors with sub-micromolar IC50 values. Beyond meeting the goals of discovering potent 3kPZS pheromone inhibitors and pioneering the use of computer-aided drug discovery for invasive species control, this project aims to advance other researchers’ success in ligand discovery by making the Screenlamp software publicly available.

## Methods

### Driving structure-activity hypothesis development by structural modeling of 3kPZS-receptor interactions

Of the available GPCR crystal structures in the Protein Data Bank [47], nociceptin, adenosine, and β1-drenergic receptors are all structural homologs [48] based on pairwise identity of 24-27% covering most of the 330-residue SLOR1 sequence [49]. SLOR1 is most similar to the β1-adrenergic receptor, based on evaluation of their sequence similarity in the extracellular loops and the inter-helical cleft comprising the orthosteric (activating ligand) binding site, the absence of non-helical insertions within transmembrane helices, and the conservation of motifs, including the E/DRY ionic lock motif in helix 3, which interacts with acidic residues in helix 6 in the inactive state of class A GPCRs [50].

A homology-based structural model of SLOR1 was constructed using the crystal structure of avian β1-adrenergic receptor as a template (PDB code: 2vt4 [51]), by using ModWeb Modeller version SVN.r972 [52]. The protein backbone structures of other related class A GPCRs with their bound ligands, such as rhodopsin, adenosine (A2A), and β1-adrenergic receptors were overlaid with the SLOR1 model to define the orthosteric binding region in SLOR1. All favorable-energy conformations of 3kPZS, generated via OpenEye OMEGA [53, 54] (v 2.4.1; https://www.eyesopen.com; OpenEye Software, Santa Fe), were docked into the SLOR1 ligand binding cavity to predict their mode of interaction by using the SLIDE software with default settings [55].

### Development of Screenlamp, a hypothesis-based screening toolkit

To facilitate the virtual screening of millions of flexible, three-dimensional structures for ligand discovery, including 3kPZS antagonists, the Screenlamp toolkit was developed in Python. It leverages high-performance memory-buffered multi-dimensional arrays [56] and data frames [57, 58]. Screenlamp first allows selection of those molecules meeting specific physicochemical or spatial properties, such as the presence of two functional groups within a certain distance. Screenlamp then interfaces with robust tools that are freely available to academic researchers to assign partial charges to ligand atoms, sample energetically favorable 3D conformers, and generate 3D overlays (OpenEye Scientific Software, Santa Fe, NM; http://www.eyesopen.com): the OpenEye molcharge utility in QUACPAC for assigning partial charges [59, 60], OMEGA [53, 54] for conformer generation, and ROCS [10, 61] for 3D molecular overlays with a reference molecule (for example, 3kPZS). The modules and tasks that can be performed within Screenlamp are summarized in Fig. 1. While an early internal version relied on an SQL database [62] for recordkeeping and an HDF5 database [63] for storing 3D coordinates of molecules, the application program interface has been simplified and computationally accelerated. Screenlamp works efficiently without SQL or HDF5 and can be applied to any molecular database organized as multi-MOL2 (3D) formatted files.

**Fig. 1.**
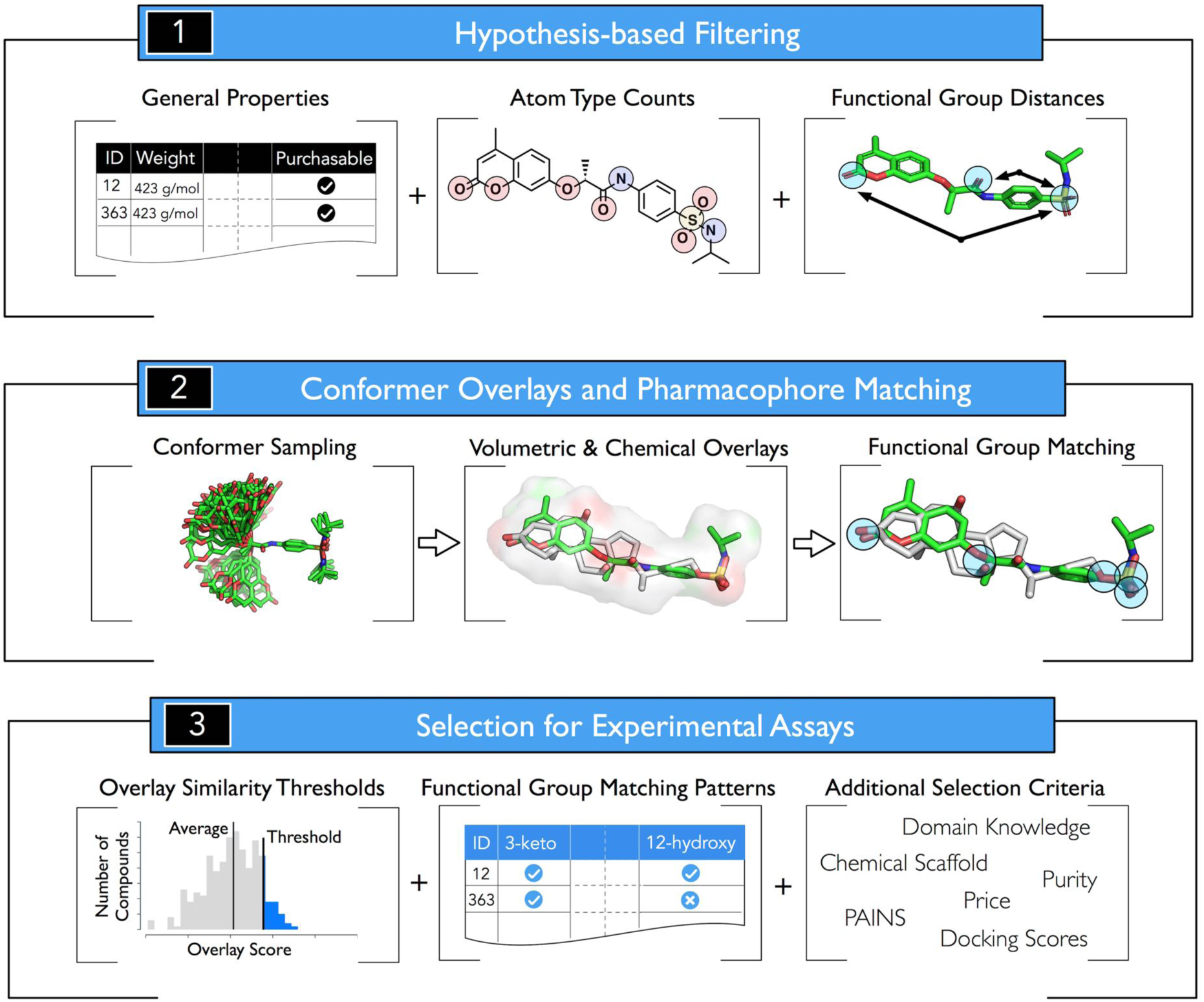
Summary of the tools provided or augmented by Screenlamp. (1) Filtering tasks that can be performed within Screenlamp to meet hypothesis-driven criteria and retrieve the structures of a subset of candidate molecules. (2) Once flexible conformers of the candidate database molecules have been sampled and overlaid with the reference molecule (for example, by using OMEGA and ROCS from OpenEye), Screenlamp can identify functional group matches in the pairwise overlays to discover functional group mimics of a reference molecule. (3) Based on the information that is available from the 3D overlays and functional group matching, as well as user-specified selection criterion, molecules are ranked for experimental testing.

The modules in the Screenlamp toolkit (Fig. 1) allow researchers to rearrange and recombine subsets of filtering, alignment, and scoring steps in a pipeline that meets their own hypothesis-driven selection criteria. For instance, a module within Screenlamp allows users to select subsets of molecules based on properties such as molecular weight, number of hydrogen bond acceptors and donors, number of rotatable bonds, or any other property data given in column format in a text file such as the property files (for example, *_prop.xls) which can be downloaded from the ZINC commercially available compound database [26]. The use of molecular property – molecular weight, number of freely rotatable bonds, etc. – can also be obtained by using open-source chemoinformatics tools such as RDKit (http://www.rdkit.org), and are optional for use in Screenlamp. Additional Screenlamp modules are available for filtering, such as selecting only those molecules that contain functional groups of interest, or optionally, functional groups in a particular spatial arrangement. Based on the 3D alignments, Screenlamp provides a module that can generate fingerprints representing the presence or absence of spatial matching between the database entries and a series of 3D functional groups in the reference molecule. These molecular fingerprints and functional group matching patterns can further be used for exploratory data analysis or machine learning-based predictive modeling of structure-activity relationships [64, 65]. Along with volumetric and electrostatic scores provided by the overlay tool, and filtering based on molecular properties, the user can then test hypotheses about the biological importance of the features he specifies by using Screenlamp to identify a matching set of compounds and procuring them for biological assays.

Once the user has selected a subset of molecules according to the current hypothesis to be tested, expressed as a set of criteria on presence or absence of certain atoms or properties, the corresponding structures are sent for conformer generation and 3D alignment with the known ligand reference molecule (typically, a known inhibitor, agonist, or substrate). The following sections provide details on how a typical workflow was implemented, in this case for the discovery of potent mimics of 3kPZS as pheromone antagonists. The Screenlamp software and full documentation are available to download from GitHub (https://github.com/psa-lab/screenlamp).

### Preparation of millions of drug-like molecules for ligand-based screening

The 3D coordinate files, in Tripos MOL2 format, of 12.3 million molecules were downloaded from ZINC12 [26] using the “drugs now” criteria (compounds with drug-like properties, available off-the-shelf). They were processed as illustrated in Fig. 1 according to the hypothesis criteria, which are summarized in the section, Hypothesis-based molecular filtering, below. Additional screening data sets of antagonist candidates were prepared, as described below, to enable the testing of close analogs of 3kPZS and known ligands of GPCRs.

#### Combinatorial analog dataset

Isomeric SMILES string (simplified molecular-input line-entry system) structural representations [66] of 332 close variants of 3kPZS were created by sampling different combinations of alternative functional groups at the 3, 7, and 12 positions in 3kPZS (Fig. 2) and different configurations (*5-α* planar or *5-β* bent relationship between the A and B rings) of the steroid ring system. These SMILES representations were used as search queries in SciFinder (http://www.cas.org/products/scifinder) to identify purchasable compounds that exactly (or nearly exactly, showing ≥99 percent similarity) match the 332 analogs. Chemical Abstract Service Registry (CAS Registry; http://www.cas.org/content/chemical-substances/faqs) identifiers were found for 84 commercially available molecules (Supplementary Material 1, Table S1). The corresponding SMILES strings were translated into 3D structures by using OpenEye QUACPAC/molcharge v1.6.3.1 (OpenEye Scientific Software, Santa Fe, NM; http://www.eyesopen.com) with the AM1BCC [60] force field for partial charge assignment. Then, OpenEye OMEGA was used to generate low-energy conformers of these molecules for virtual screening.

**Fig. 2.**
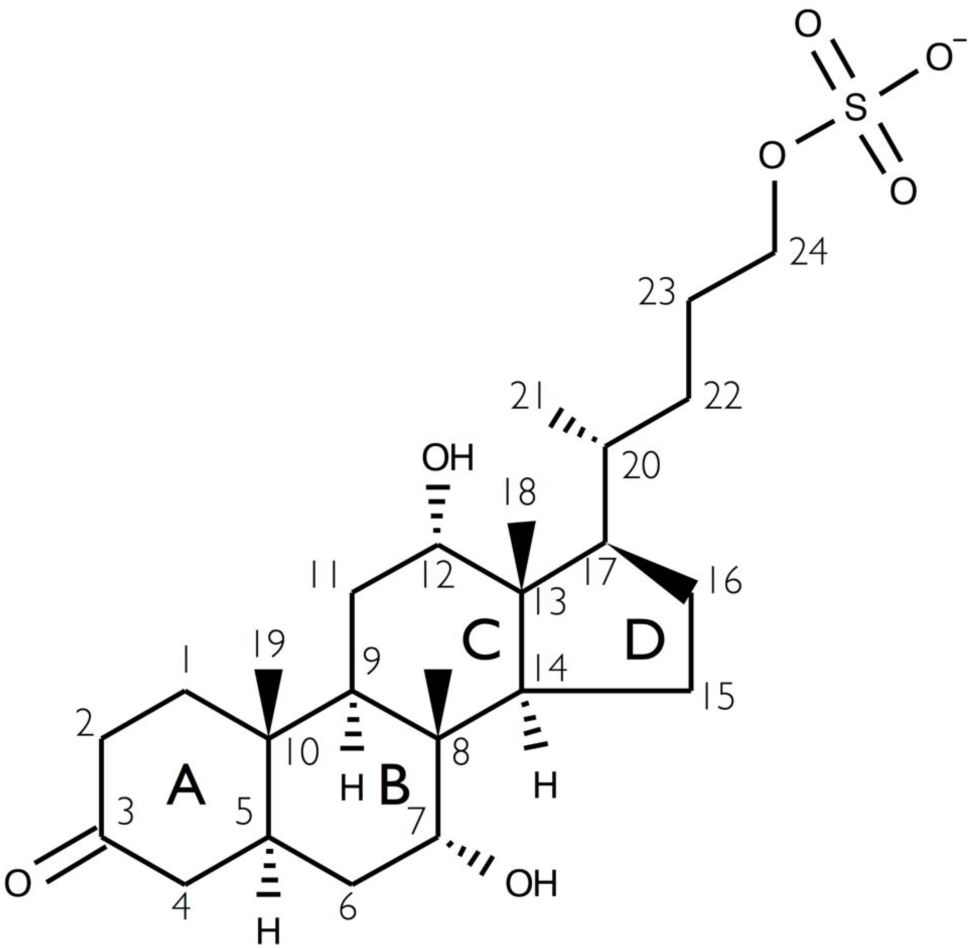
The molecular structure of 3kPZS, showing canonical atom numbers and ring labels for the steroid ring system and tail (positions C-20 to C-24).

#### CAS Registry steroids

The ZINC database covers many, but not all, vendors of small organic molecules; thus, the CAS Registry of 91 million compounds (https://www.cas.org/content/chemical-substances) was searched with SciFinder Scholar (Chemical Abstracts Service, Columbus, OH) for all commercially available steroid molecules that were not already present in the ZINC database. Using SciFinder (which limits the number of molecules that can be processed at a time to 100), batches of CAS Registry steroid structures were exported and processed into SMILES strings using CACTUS (http://cactus.nci.nih.gov). Three-dimensional structures were created from the SMILES strings as described for the combinatorial analog dataset above, resulting in 2,995 additional steroids for screening (Supplementary Material 2, Table S2).

#### GPCR Ligand Library (GLL)

The GLL database consists of approximately 24,000 known ligands for 147 GPCRs [67] (http://cavasotto-lab.net/Databases/GDD/). To prepare this database for our virtual screening pipeline, partial charges were added to the existing 3D structures of these molecules using OpenEye QUACPAC/molcharge with the AM1BCC force field, and low-energy conformations were generated using OpenEye OMEGA with default settings.

### Identification of incorrect steroid substructures in molecular databases

In version 12 of ZINC (http://zinc.docking.org), if a vendor did not provide complete stereochemistry information for chiral centers in a steroid molecule, up to four different stereoisomeric structures were automatically provided by ZINC, each with a separate ID. However, at most one of those four structures had a valid steroid configuration (with a *5-α* planar or *5-β* ring structure and 18- and 19-methyl group orientations as shown in Fig. 2). Thus, we developed a custom steroid checking tool using the OpenEye OEChem toolkit by comparing each molecule with an isomeric SMILES representation of the canonical steroid core atom connectivity and chirality, to filter out invalid steroid configurations. This steroid checker is included in Screenlamp and has recently been implemented in ZINC, by coordination with the developers at UCSF.

#### Step 1: Hypothesis-based molecular filtering

The Screenlamp toolkit provides a user-friendly interface to efficiently select those molecular structures relevant to a given screening hypothesis or objective. For instance, the first step in the 3kPZS inhibitor screening (Fig. 3) selected those drug-like molecules listed as commercially available by either ZINC or CAS. Drug-like properties were defined as satisfying Lipinski’s rule of 5 [68], with an additional rotatable bond criterion to filter out highly flexible molecules because their significant loss of entropy upon protein binding detracts from the ΔG_binding_ between receptor and ligand. The drug-like criteria used were: (1) molecular weight between 150 and 500 g/mol; (2) octanol-water partition coefficient less than or equal to 5; (3) 5 or fewer hydrogen bond donors and 10 or fewer hydrogen bond acceptors; (4) polar surface area less than 150 Å^2^; and (5) fewer than 8 rotatable bonds. In addition, the filtering query excluded all molecules that were flagged as invalid steroids.

**Fig. 3.**
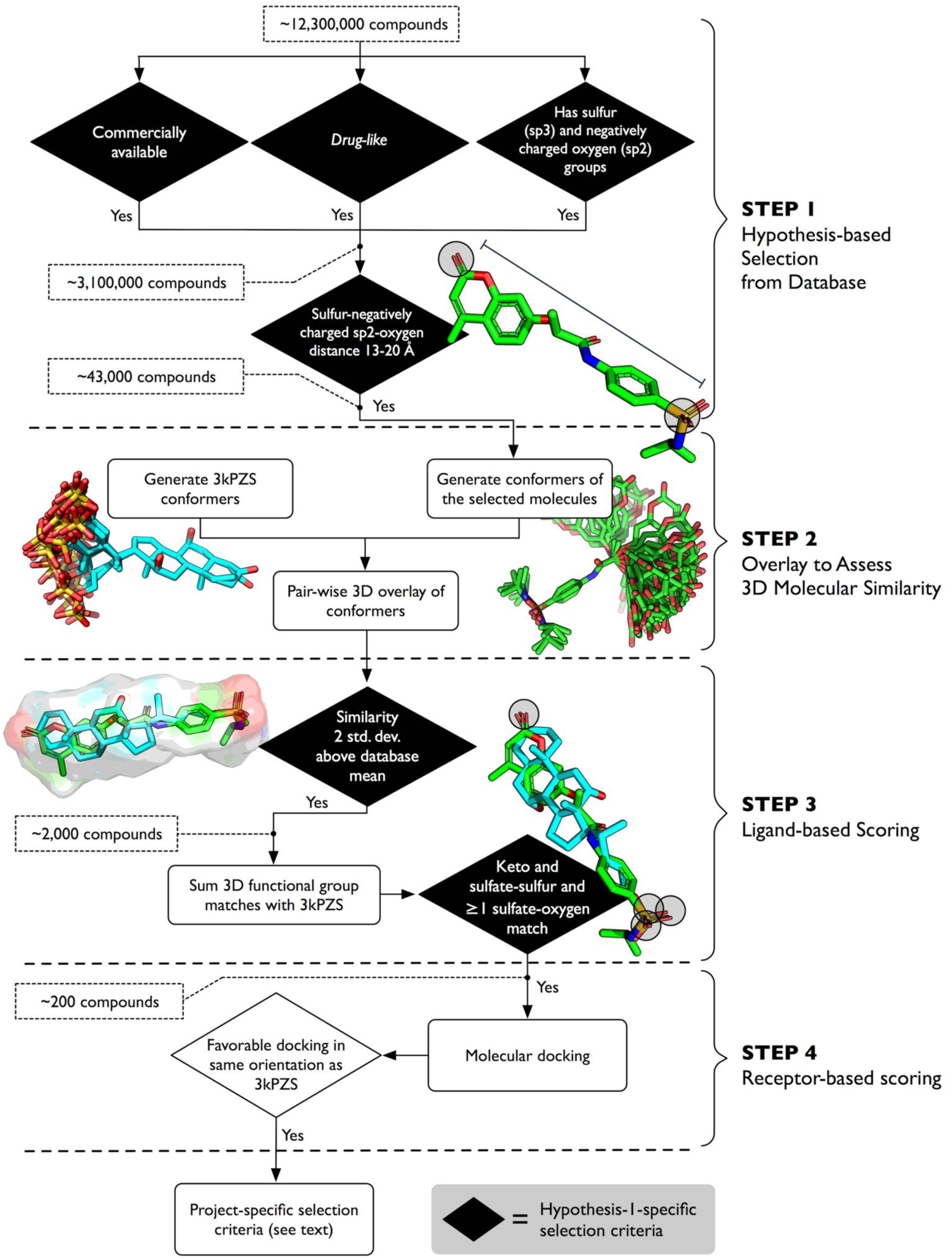
Using Screenlamp to identify compounds to test the hypothesis that compounds with negatively charged sulfate and sp2-hybridized oxygen groups matching the 24-sulfate-3-keto oxygen distance in 3kPZS will mimic 3kPZS and block its binding. Note that the mean and standard deviation in TanimotoCombo score used in Step 3 to filter candidate molecules was derived from the score distribution of the single conformers from the entire ZINC drug-like database overlayed with the panel of 3kPZS conformers (rather than from the set of 43,000 molecules at the end of Step 1); see Supplementary Materials 3 Fig. S1 for the distribution of ZINC database TanimotoConformer scores following overlay with 3kPZS.

#### Step 2: Sampling favorable molecular conformations

Low-energy conformations of the reference molecule 3kPZS were generated by sampling rotatable bond torsions and flexible ring systems with OpenEye OMEGA [53, 54] v 2.4.1, using its default settings. Forty-eight favorable 3D conformers in a somewhat extended rather than folded conformation, required to fit within the ligand-binding site of the SLOR1 structure (Fig. 1), were kept (Fig. 3); specifically, the distance between the 3-keto group and the sulfur atom in the sulfate group in these conformers was 13-20 Å. Up to 200 favorable (low) energy conformations, following clustering by OMEGA to identify distinct conformations, were retained for each of the database molecules selected by the Screenlamp filtering steps. OMEGA uses a modified version of the Merck molecular force field (MMFF94) to measure the internal energy of a given conformation and removes conformers with internal clashes. Based on the lowest energy conformer, up to 199 additional conformers are selected by OMEGA if their energy scores are less than 10 kcal/mol higher than the lowest energy conformer and they have a pair-wise atom-position RMSD value of at least 0.5 angstroms when overlayed with every other conformer in the set.

### Generation of overlays to compare molecular shape and charge distribution with a known ligand

In addition to property and pharmacophore-based filtering, Screenlamp invokes the ROCS [10] software to generate 3D molecular overlays to evaluate similarity in volumetric and chemistry (or “color” [69]). In particular, the color component evaluates whether two atoms are attractive (same color atom type) or repulsive (opposite color atom types), including weighting terms that measure the strength of the interaction. The color types include H-bond donor, H-bond acceptor, anion, cation, hydrophobic, and component of a ring. Using OpenEye ROCS v 2.4.6, 48 low-energy conformers of the 3kPZS reference molecule were overlaid with up to 200 conformers for each of the selected database molecules. The 3D overlays were ranked by the TanimotoCombo metric, which consists of equally contributing components that assess the degree of volumetric (shape) and atom type (color) similarity upon alignment. The TanimotoCombo metric requires perfect match between all parts of two molecules (rather than an exact substructure match) to achieve a perfect score, which ranges between 0 (no overlap/similarity) and 2 (perfect overlap). For each database entry, only the configuration of the best-overlaid pair of conformers between 3kPZS and the database molecule was saved.

#### Step 3: Ligand-based scoring

Molecules with a similarity score two standard deviations above the mean, showing a high degree of similarity to the 3kPZS reference molecule, were considered as potential 3kPZS mimics and evaluated for functional group matches with 3kPZS. Here, the similarity score threshold was computed from overlaying the low-energy 3kPZS conformers with the single-conformer ZINC drug-like database, which resulted in a mean TanimotoCombo score of 0.65 with a standard deviation of 0.083 (Supplementary Material 3, Fig. S1). 307,517 ZINC drug-like molecules have a TanimotoCombo score of 0.82 or greater (0.65 plus two standard deviations) when overlayed with the 3kPZS multi-conformer query. This TanimotoCombo threshold was relaxed for some screening hypotheses to allow selection of more compounds with favorable matches according to additional criteria, as detailed in the section, Hypothesis-driven selection of ligand candidates, below. Following overlay by ROCS, functional group matches were identified in the database molecule based on the presence or absence of an atom with atom type, atomic charge, and hybridization matching the following (numbered according to the positions in 3kPZS): 3-keto, 3-hydroxyl, 7-hydroxyl, and 12-hydroxyl oxygens; 18- and 19-methyl groups; sulfate ester oxygen and three sulfate terminal oxygens (Fig. 2). In each case, an atom (functional group component) in the database molecule was considered to match an overlayed atom in 3kPZS if its atom type, atomic charge, and hybridization matched, and the two atom centers were within 1.3 Å in the highest-scoring ROCS overlay of the database molecule with 3kPZS.

### Docking the highest-ranking compounds with the SLOR1 structural model to assess goodness of fit

For inhibitor candidates prioritized by two of the hypotheses (3 and 8, described in the following section), flexible docking was performed by using SLIDE v 3.4 with default settings [55] to compare the mode of interaction of a given ligand candidate with 3kPZS docked into SLOR1. It was noted whether each candidate docked with a favorable SLIDE-predicted ΔG_binding_ and whether it could form a salt bridge with His110 in SLOR1, similar to that observed for the 3kPZS sulfate tail, as described in the Results. His110 proximity to the sulfate or negatively charged group in docked ligands selected by hypotheses 3 and 8 was used to define whether the fundamental orientation of the ligand candidate in the narrow binding cleft allowed formation of this favorable electrostatic interaction hypothesized to be essential for pheromone-like binding, or alternatively whether its sulfate or sulfate-like group was oriented outward, towards the extracellular opening of the cleft.

### Assays to measure inhibition of olfactory response of 3kPZS

Electro-olfactogram assays (EOGs) are commonly used to measure *in vivo* olfactory responses to environmental stimuli in vertebrates [70]. EOGs record the sum of action potentials (the field potential) generated upon the activation of olfactory receptors (predominantly GPCRs) in the olfactory epithelia upon exposure to an odorant. The sea lamprey EOG assays were conducted following a standard protocol described in Brant *et al.* 2016 [71]. Adult sea lamprey were anesthetized with 100 mg/L of 3-aminobenzoic acid ethyl ester (MS222, Sigma-Aldrich Chemical Co.) and injected with 3 mg/kg gallamine triethiodide (Sigma-Aldrich Chemical Co.). Then, the gills were exposed to a continuous flow of aerated water with 50 mg/L MS-222 throughout the experiments. All tested compounds were delivered directly to the olfactory rosette using a small capillary tube. Water used in the EOGs was charcoal filtered fresh water. At the beginning of each experiment, and after each compound was tested, the olfactory rosette was flushed with charcoal filtered fresh water for 2 minutes before the responses to 3kPZS (10^-6^ M) and _L_-arginine (10^-5^ M), a strong sea lamprey olfactant that is not a pheromone, were recorded. To test the effect of the inhibitor candidates on the olfactory detection of 3kPZS, the olfactory rosette was continuously exposed to a 10^-6^ M solution of the candidate for 2 minutes. The responses were measured for a mixture of 10^-6^ M 3kPZS and 10^-6^ M of the inhibitor candidate, and also for a mixture of 10^-5^ M _L_-arginine and 10^-6^ M of the candidate. The two mixtures were recorded for 4 seconds each. The percent reduction of the 3kPZS olfactory response was calculated as: 1 – ([response of the 3kPZS and candidate mixture – response to blank charcoal filtered fresh water] / [3kPZS response before inhibitor candidate – response to blank charcoal filtered fresh water]) × 100. Compounds that reduced the EOG response to 1 × 10^-6^ M 3kPZS by 50% or more when combined with 3kPZS in equimolar concentration are described as having 1 micromolar or lower IC50 values. If the compound either inhibited or acted as an agonist of the 3kPZS receptor (competing for the receptor binding site with 3kPZS), a reduction of the 3kPZS signal would be observed. The response of a mixture of L-arginine and the inhibitor candidate was recorded, and the percent reduction of the L-arginine response was calculated to ensure that the inhibitor candidate had a specific effect in the 3kPZS receptor. L-Arginine is a common stimulant of the olfactory epithelium of sea lamprey that does not compete with the 3kPZS signal transduction pathway or receptor [72]. Recordings for each compound were repeated two to five times, and the reported signal reduction was computed as the average signal reduction among the replicates.

### Graphics tools

Molecular graphics and renderings were produced using MacPyMOL v1.8.2.2 [73]. Data plots were generated using matplotlib v2.0.0 [74], and diagrams were drawn using vector graphics software OmniGraffle v7.3.1, Affinity Designer v1.5.5, and Autodesk Graphic v3.0.1. All images were exported into bitmap format using macOS Preview v9.0.

## Results and Discussion

### Structural model for interactions between 3kPZS and SLOR1

A 3D atomic structure of SLOR1 was built using the ModWeb interface to MODELLER and energy-minimized with CHARMM, as described in the Methods. MODELLER computed 25.5% sequence identity between SLOR1 and the β1-adrenergic receptor template (PDB entry 2vt4), with a highly significant expectation value of 6.4e^-11^. Structural evaluation by PROCHECK showed 94% of the SLOR1 residues to have main-chain dihedral values in the most-favored region, comparable to high-resolution crystal structures, and overall favorable stereochemistry, as measured by the PROCHECK G-factor (Supplementary Materials 4). SwissModel Workspace evaluation tools indicated that the model has favorable all-atom contacts [75] and a favorable overall energy [76] similar to those in the 2vt4 template structure (Supplementary Materials 5 and 6). The structural model of SLOR1 and generation of flexible conformers starting with the 3D structure of 3kPZS determined by NMR and mass spectrometry [27] enabled prediction of their interaction by docking. The most favorable docking mode predicted by SLIDE, with a predicted Δ G_binding_ of -9.0 kcal/mol, showed the sulfate tail binding deeply in the orthosteric cleft near His110, surrounded by the transmembrane helices and open to the extracellular space (Supplementary Materials 7, 8, and 9). The planar steroid binds in this cleft almost parallel to the transmembrane helical axes, with the specificity-determining 3-keto group pointing towards the solvent-exposed extracellular loops (Fig. 4).

**Fig. 4.**
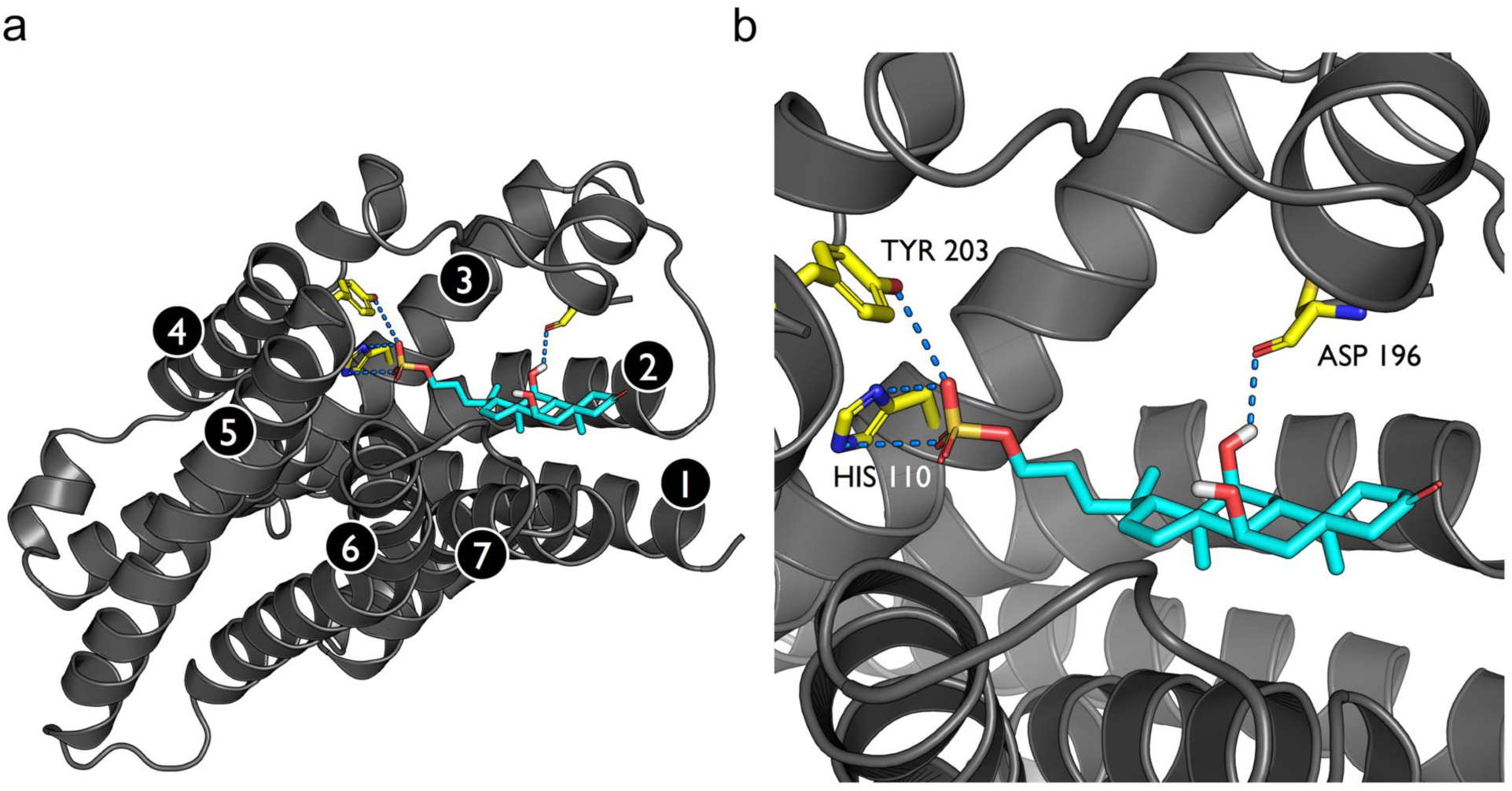
The backbone of the SLOR1 homology model is shown in gray ribbons with side-chain carbon atoms in yellow. 3kPZS carbon atoms are shown in cyan, with oxygen atoms in red, nitrogen atoms in blue, the sulfate sulfur in yellow, and hydrogen atoms in white. Hydrogen bonds between SLOR1 and 3kPZS are drawn as blue dashed lines. (a) Full view of the SLOR1-3kPZS complex, with GPCR transmembrane helices enumerated 1-7 from N-to C-terminus. (b) Close-up of key polar side chains forming intermolecular hydrogen bonds and salt bridges between SLOR1 and 3kPZS, as determined by SLIDE docking.

The main sulfate-binding residue, His110, is 10 Å above a regulatory sodium site elucidated in the high-resolution adenosine receptor structure and thought to occur in many class A GPCRs (PDB entry 4eiy [77]). Most of the sodium-ligating side chains are identical or similar in side-chain chemistry in SLOR1 (Fig. 5). The strongly attractive salt bridge between the 3kPZS sulfate group with both side chain nitrogen atoms on neighboring His110, reinforced by a hydrogen bond with Tyr203 and through-space electrostatic attraction with the postulated buried sodium ion [78], help explain the sensitivity of SLOR1 to 3kPZS in low nanomolar concentration. The di-methylated face of the steroid system in 3kPZS (Fig. 2 and Fig. 4) is predicted to bind to a highly hydrophobic wall in the SLOR1 cleft, comprised of hydrocarbon side chain groups from Phe87, Met106, Leu109, His110, Asp196, Pro277, Tyr280, and Thr284. The 12-hydroxyl group on the opposite face of the steroid ring hydrogen-bonds with the Cys194 main-chain oxygen. This mode of interaction is supported by a very similar cholate binding prediction for SLOR1 from CholMine [79] (http://cholmine.bmb.msu.edu), in which Phe87, Leu109, and His110 also form major interactions with the dimethylated edge of the steroid ring system. This comparison between the binding mode predicted by CholMine for cholate and the interaction of 3kPZS predicted by SLIDE docking for SLOR1 can be visualized with the PDB and PyMOL graphics files available in Supplementary Materials files 10, 11, 12, and 13. The position of the 3-keto group at the solvent interface, not directly contacting SLOR1, suggests it interacts with the thirty N-terminal residues of SLOR1 that are absent from the model due to lack of homology with any PDB structure. Structures of ligand-interacting lid peptides in class A GPCRs (which includes the CDYLVVLFL sequence in SLOR1) are highly individualized according to receptor type [78]. Together, the interactions predicted between the 3kPZS and SLOR1 structures informed the development of several screening hypotheses, as described below.

**Fig. 5.**
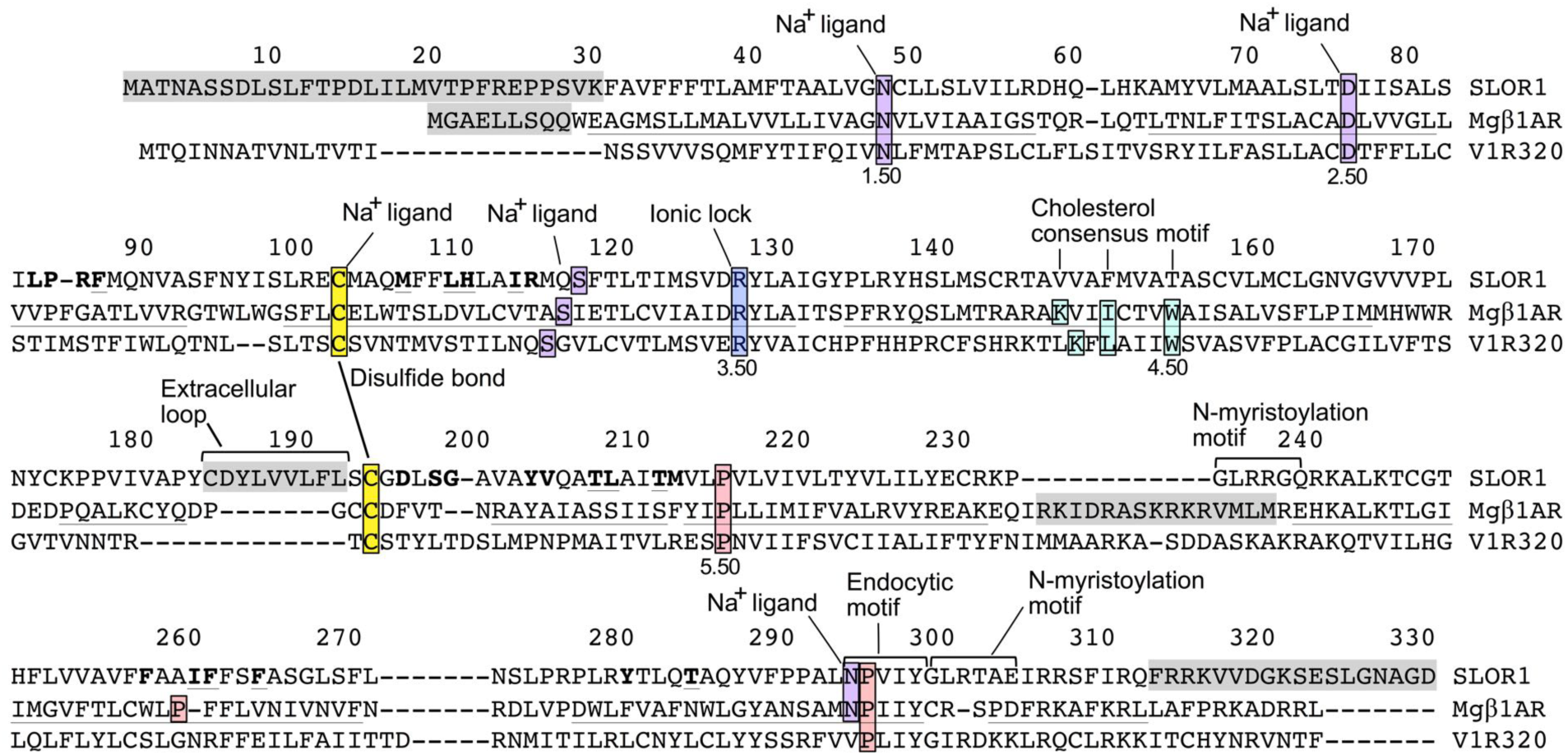
Alignment of the sea lamprey SLOR1 sequence with the closest GPCR of known 3D structure, β1-adrenergic receptor from *Meleagris gallopavo* (turkey), labelled Mgβ1AR. The V1R320 receptor [36], another class A sea lamprey GPCR activated by 3kPZS, is shown for comparison. Highlighted residues form a disulfide bond (yellow) and cholesterol (aqua) and sodium ion (purple) binding sites, as mentioned in the text. Putative N-myristoylation and endocytic binding motifs are labeled and indicated by brackets above the sequence. The most highly conserved residue in each helix (highlighted in pink, if not already highlighted according to one of the above roles) is labeled by its Ballesteros-Weinstein GPCR sequence number X.50, where X indicates the transmembrane helix number and 50 is the position number assigned to the most conserved residue in that helix across all GPCRs [86]. Residue numbers above the alignment correspond to the full SLOR1 sequence and are right-justified to align with the corresponding single-letter amino acid code Boldface indicates predicted ligand binding site residues in SLOR1, based on occurring within 5Å of the retinal ligand in rhodopsin (PDB entry 2z73) or cyanopindolol in the β1-adrenergic receptor structure (PDB entry 2vt4), following their structural superposition on SLOR1 by DaliLite (http://ekhidna.biocenter.helsinki.fi/dali_lite). Gray highlighting indicates residues with no structural model in SLOR1 due to low homology with the β1-adrenergic receptor (Mgβ1AR) or absence of crystallographic coordinates in this region of the Mgβ1AR crystal structure (PDB entry 2vt4). Underlined residues form the transmembrane helices in Mgβ1AR, based on DSSP main-chain hydrogen-bonding analysis provided by the PDB [87].

### Hypothesis-driven selection of ligand candidates

Upon screening the compounds from the ZINC, CAS and GLL databases to identify those with significant ROCS TanimotoCombo scores and functional group matches with 3kPZS as tabulated by Screenlamp (see Methods), top-scoring compounds were selected for electro-olfactogram (EOG) assays to directly assess their ability to reduce the sea lamprey olfactory response of 3kPZS, according to the following hypotheses. Several of the hypotheses involve the presence of oxygen in a spatial position equivalent to the 3-keto oxygen in 3kPZS and negatively charged oxygen in positions equivalent to the terminal sulfate oxygens in 3kPZS. The focus on these functional groups is based on prior results indicating that both 3-O and sulfated tail groups were associated with high olfactory potency [80] and were present in a few other steroid compounds known to elicit 3kPZS-like activity [81]. Specific hypotheses that were tested:

1. *Compounds matching the 3-keto and one or more sulfate oxygens in extended conformations of 3kPZS*, with TanimotoCombo score greater than 2 standard deviations above the mean. This tests the hypothesis that high overall shape and electrostatic similarity and matching the 3-keto and sulfate groups are sufficient to mimic and block 3kPZS activity.
2. *Similar to the above, compounds with 3-hydroxyl groups spatially matching the 3-keto moiety in 3kPZS* were selected to test the hypothesis that 3-hydroxyl containing compounds can block 3kPZS olfaction. Additional criteria for this set: ROCS similarity score to 3kPZS of 0.8 or above, high ROCS ColorTanimoto score (0.25 or above), matching a sulfate terminal oxygen and at least one of the other functional groups in 3kPZS (sulfate oxygen, hydroxyl or steroid methyl substituents), and docking with the sulfate group proximal to His110 in SLOR1 with a favorable predicted Δ G_binding_ (< -7 kcal/mol).
3. *All compounds with a planar steroid ring system and alpha configuration of hydroxyl groups matching 3kPZS* (rather than equatorial configuration), 3-keto and sulfate oxygen matches, and ROCS TanimotoCombo scores greater than 0.65. This set tests the hypothesis that close steroidal analogs matching the oxygen-containing groups in 3kPZS will mimic 3kPZS activity. The emphasis on planar (5-*α*) steroids derives from the fact that sea lamprey are the only fish that synthesize planar steroids with sulfated tails [82], and these features are expected to be species-selective olfactory cues. In fact, 5-*β* steroid relatives of 3kPZS are far less potent [81].
4. *Phosphate or sulfate tail analogs. Aliphatic chains with at least 3 methyl(ene) groups terminating in a phosphate or sulfate group* were identified by the ZINC search tool, to test whether mimicking the sulfate tail moiety of 3kPZS alone is sufficient to block 3kPZS olfaction, and whether a phosphate group can mimic the sulfate group.
5. *Compounds with a high degree of shape/electrostatic match with the C and D steroid rings and sulfated aliphatic tail structure in PAMS-24*, another sea lamprey pheromone identified by the Li lab [83] (Supplementary Material 3, Fig. S2). This region corresponds to atoms 8, 9, and 11-24 plus the sulfate group (Fig. 2), with the addition of an isopropyl group at C-24. This set tests whether compounds matching a pheromone tail fragment inhibit the 3kPZS response.
6. *5-β steroid structures with at least 2 sulfate oxygen matches with 3kPZS or at least 5 functional group (oxygen and methyl) matches* were chosen to test whether bent rather than planar steroids can block 3kPZS olfaction. (None of the compounds could match both the 3-keto and sulfate tail, due to the bent geometry of the 5-β steroid ring system when overlaid with ROCS to match all atoms.) Prior work on 5-β steroids tested their ability to act as 3kPZS agonists rather than inhibitors [81].
7. *Compounds with highly negative sulfate oxygen-matching atoms* (with charges at least 0.3 units more negative than the sulfate oxygen charge in 3kPZS) were selected, testing the hypothesis that strongly negatively charged groups can form stronger interactions with SLOR1 (e.g., salt bridge with His110) and outcompete 3kPZS for binding.
8. *Compounds with negatively charged, non-oxygen-containing tails*, testing whether other negative groups can block 3kPZS binding. These compounds contained a negatively charged atom other than oxygen in one of the sulfate oxygen positions, a high ROCS ShapeTanimoto score for similarity to 3kPZS (0.8 or above) and favorable ROCS ColorTanimoto value (0.25 or above). Further filtering criteria included matching the 3-hydroxyl group and a sulfate terminal oxygen, at least one of the other functional groups (sulfate oxygen, hydroxyl or steroid methyl substituents), and docking with the sulfate group close to His110, deep in the SLOR1 binding site, with a docking score < -7 kcal/mol.
9. *Epoxide-containing steroids.* Epoxide functional groups are labile, tending to spring open due to bond strain and then reacting with nearby protein groups. Previous research [84] indicated that epoxide cross-links can be site-specific, preferring histidine side chains. 3kPZS-like, epoxide-containing steroids were tested because cross-linking with the active-site His110 in SLOR1 could result in very strong inhibition of the 3kPZS receptor. Epoxide opening or cross-linking from the equivalent of the 3-oxygen position in 3kPZS could also create an antenna-like group, potentially making favorable interactions with the collar of the binding site as has been found for other GPCR ligands [85].
10. *Taurine tail-containing steroids with significant overall chemical similarity to 3kPZS.* Taurolithocholic acid was observed to significantly block the olfactory response of sea lamprey to 3kPZS; in Fig. 7, taurolithocholic acid is compound #3 and was observed to block 65% of the 3kPZS response. Other compounds with taurine tails and overall good matches to 3kPZS could also block its binding to SLOR1 and were identified as candidates for testing.
11. *CAS steroids with high volumetric and electrostatic similarity to 3kPZS.* The top 25 steroids from the CAS Registry, ranked by volumetric and electrostatic similarity to 3kPZS in ROCS overlays, were selected for experimental assays. These compounds are highly similar to 3kPZS while not being biased by prior knowledge of activity determinants, by which we mean chemical groups or pharmacophores involved in receptor binding.
12. *Compounds known to be bioactive in complex with the β-adrenergic receptor*, the GPCR of known structure with highest binding site sequence similarity to SLOR1. Three molecules known to be active versus β1-adrenergic receptor were selected for assaying: carvedilol (agonist; ZINC01530579), atenolol (selective antagonist; ZINC00014007), and dobutamine (partial agonist; ZINC00003911).

**Fig. 7.**
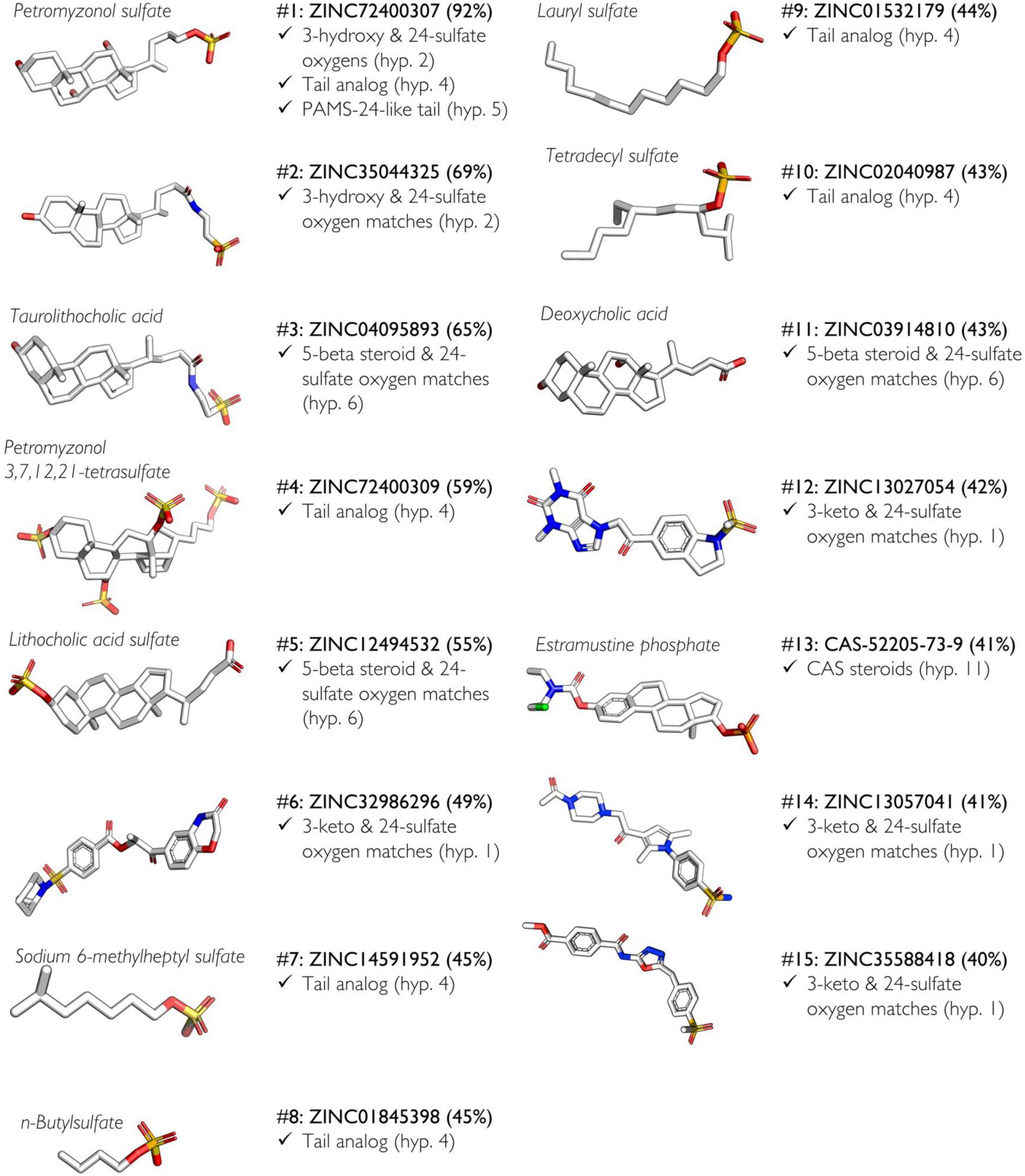
3D structures of the 15 most active molecules from screening the ZINC drug-like dataset, the combinatorial analog dataset, CAS registry steroids, and the GPCR Ligand Library as described in the Methods section, *Preparation of millions of drug-like molecules for ligand-based screening.* The molecule structures are numbered from highest percent inhibition (#1) to lowest (#15). Accession codes in the CAS registry and ZINC databases are provided along with average percent inhibition values over two or more replicates. Hypothesis-based selection criteria are listed below the compound IDs, referencing the hypothesis descriptions given in the Results. The ZINC13057041 compound has been flagged as a potential PAIN (pan-assay interference compound) containing a functional group that leads to false-positive assay results via the server available at http://cbligand.org/PAINS/ [88].

### Screenlamp discovery of potent 3kPZS antagonists

A typical screening run for a single hypothesis, for instance, obtaining 3kPZS volumetric and pharmacophore mimics with 3-oxygen and 24-sulfate matches (Fig. 3) starting from 12 million commercially available, drug-like molecules in ZINC, was completed within a day on a standard desktop computer (2 Intel^®^ Xeon^®^ CPU E5-2620 v2 at 2.10GHz (8 cores), 16 GB DDR3 SDRAM, and 7200 RPM hard drive). Candidates from the hypothesis-based screens described in the Methods provided a set of 307 commercially available compounds, including 8 samples from different vendors for some of the 299 unique compounds. The entire set was procured and tested by EOG for the ability to reduce the 3kPZS olfactory response. Following the EOGs, the most and least active compounds (Fig. 6) were analyzed structurally to identify features that correlate with activity.

**Fig. 6.**
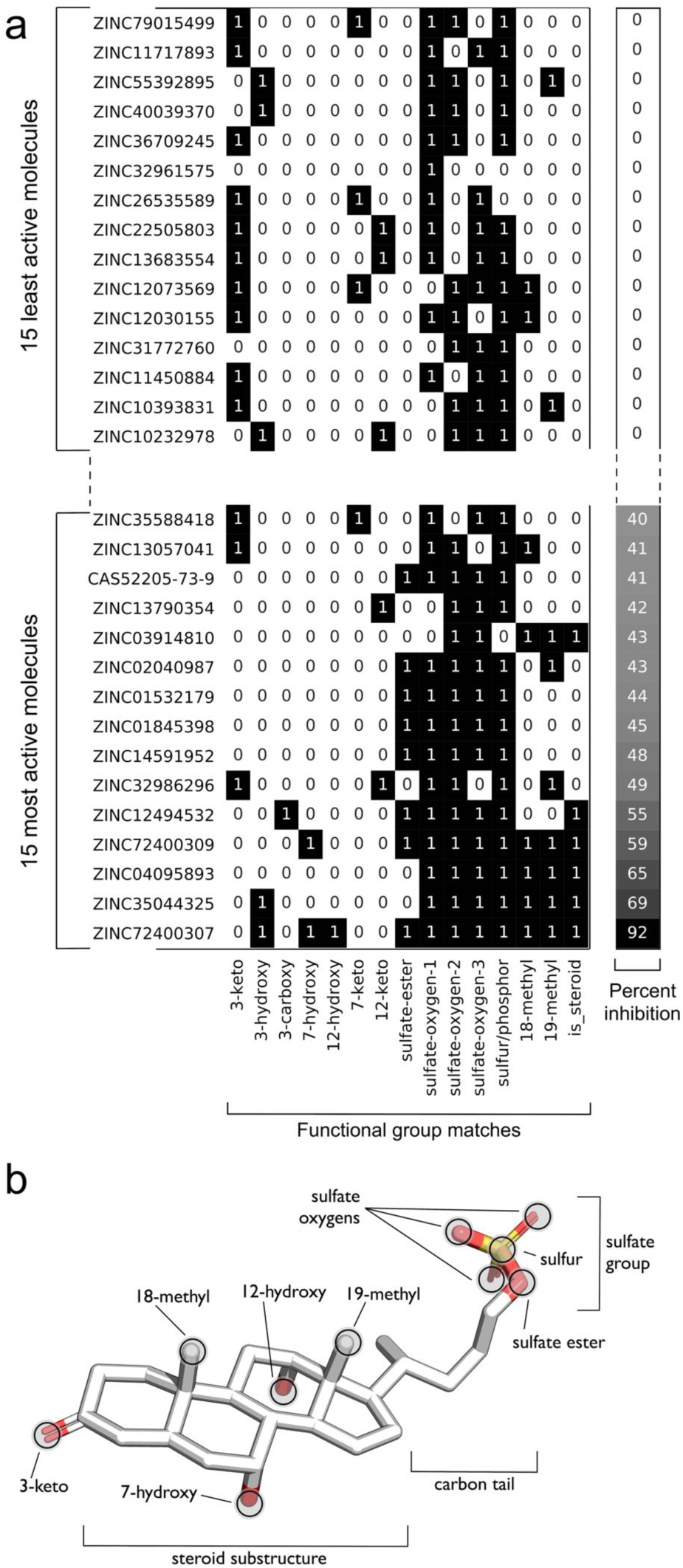
(a) Heat map showing the functional group matches of the 15 most active and 15 least active molecules when overlaid with 3kPZS. The percent inhibition was computed as the average inhibition over two or more independent electro-olfactogram assays. Heat map cells containing 1’s indicate the presence of a match and 0’s indicate the absence of a match. (b) 3D representation of an energetically favorable 3kPZS conformer with functional group labels, to aid in interpreting the heat map x-axis labels. We have developed computational protocols to identify activity discriminants from such data; see [65] and the example code on GitHub: https://github.com/psa-lab/predicting-activity-by-machine-learning.

### Structure-activity relationships of Screenlamp compounds

Six of the 15 most active compounds, which reduced the response to 3kPZS by 43-92%, were steroidal. Interestingly, most of the top 15 inhibitors other than petromyzonol sulfate (PZS; ZINC72400307; the 3-hydroxyl analog of 3kPZS), lacked 3kPZS-like hydroxyl groups in the 7- and 12-positions of the steroid ring system. The three-dimensional overlays of all 15 compounds with the best-matching conformer of 3kPZS are provided in Supplementary Materials file 14. The three most active compounds had 3-hydroxyl groups in place of the 3-keto group in 3kPZS (Fig. 7), suggesting that this group acts as a switch between agonist and inhibitor functions. These results highlight the importance of performing some screens without filtering heavily on chemical group matches, because our initial screens focused on the hypothesis that presence of a keto group that matches the 3-keto group in 3kPZS is essential for binding the 3kPZS receptor. The most active compound discovered in this work, PZS, which contains a 3-hydroxy group instead of 3-keto (Fig. 7), first arose from our screen based on hypothesis 4 (presence of a sulfate tail). The discovery that PZS was highly active, yet contained a 3-hydroxy group, drove the development of the successful hypothesis 2 screen, seeking 3-hydroxy and 24-sulfate tail matches. (Note that the hypotheses are numbered arbitrarily, not in the order that they were performed. Several of the screens were performed concurrently, while others were initiated as assay results provided new insights into chemical groups associated with activity.) For the two most active compounds (both selected by hypothesis 2), the inhibitor orientation was found to be 3kPZS-like, energetically favorable, and capable of forming a salt bridge with His110 deep in the orthosteric pocket of SLOR1. In these two compounds, PZS and taurolithocholic acid (ZINC35044325), the 3-hydroxyl groups overlapped with the 3-keto group of 3kPZS upon 3D overlay by ROCS (Fig. 6).

Other interesting structure-activity relationships were revealed by six sulfate tail analogs (hypothesis 4), matching the sulfate tail moiety of 3kPZS, appearing among the 10 most active compounds (Fig. 7). For instance, the four compounds ZINC14591952 (sodium 6-methylheptyl sulfate), ZINC01845398 (n-butylsulfate), ZINC01532179 (lauryl sulfate), and ZINC02040987 (tetradecyl sulfate) consist entirely of aliphatic hydrocarbon chains terminating in a sulfate group (Fig. 7) and were found to reduce the olfactory response of 3kPZS by 43-45%. This is a useful insight, as it indicates that matching the 3-keto oxygen and steroid ring system in 3kPZS is not absolutely essential. Sulfated alkanes like these are inexpensive compounds, though they vary in vertebrate toxicity and are likely to be less target-selective than molecules capable of making additional 3kPZS-like interactions. The trisulfated variant of PZS (ZINC72400309) is another molecule that would not have been predicted by a typical drug discovery approach, due to its high polarity and bulk relative to the reference compound, 3kPZS. However, this turns out to be one of the most effective antagonists of 3kPZS according to *in vivo* olfactory inhibition of 3kPZS response. Trisulfated PZS shows even greater promise in behavioral tests with sea lamprey in natural stream water (manuscript in preparation). The other hypotheses that were most effective, discovering moderate-activity compounds that blocked 40-65% of the olfactory response to 3kPZS, were hypothesis 1 (compounds matching the 3-keto and 24-sulfate groups in 3kPZS) and hypothesis 6 (compounds containing a 5-β steroid ring and 24-sulfate group match).

### Enrichment of active molecules through hypothesis-based filtering criteria

While shaped-based ranking methods for ligand-based virtual screening such as ROCS yield favorable results when tested for the ability to identify active molecules among a large set of decoys [10], the development of Screenlamp was driven by the need to include domain-based knowledge, such as spatial relationships between a subset of functional groups observed in pheromones, to discover compounds highly enriched in activity. The benefits of incorporating system-specific criteria when screening is underscored by considering the results of using only shape and charge scoring. Upon comparing the TanimotoCombo scores alone, based on shape and pharmacophore similarity to 3kPZS, with EOG assay values representing the percent inhibition of 3kPZS response for the 299 compounds (Supplementary Material 15, Table S3), the correlation between molecular similarity scores and percent inhibition of 3kPZS response values was low (Fig. 8), as indicated by a Pearson linear correlation coefficient value (R) of 0.07. On the other hand, the consistently accurate 3D overlays with 3kPZS provided by ROCS for the entire screening database were essential for prioritization of compounds prior to chemical group matching, and the accurate ROCS overlays also enabled correct detection of chemical group matching in Screenlamp.

**Fig. 8.**
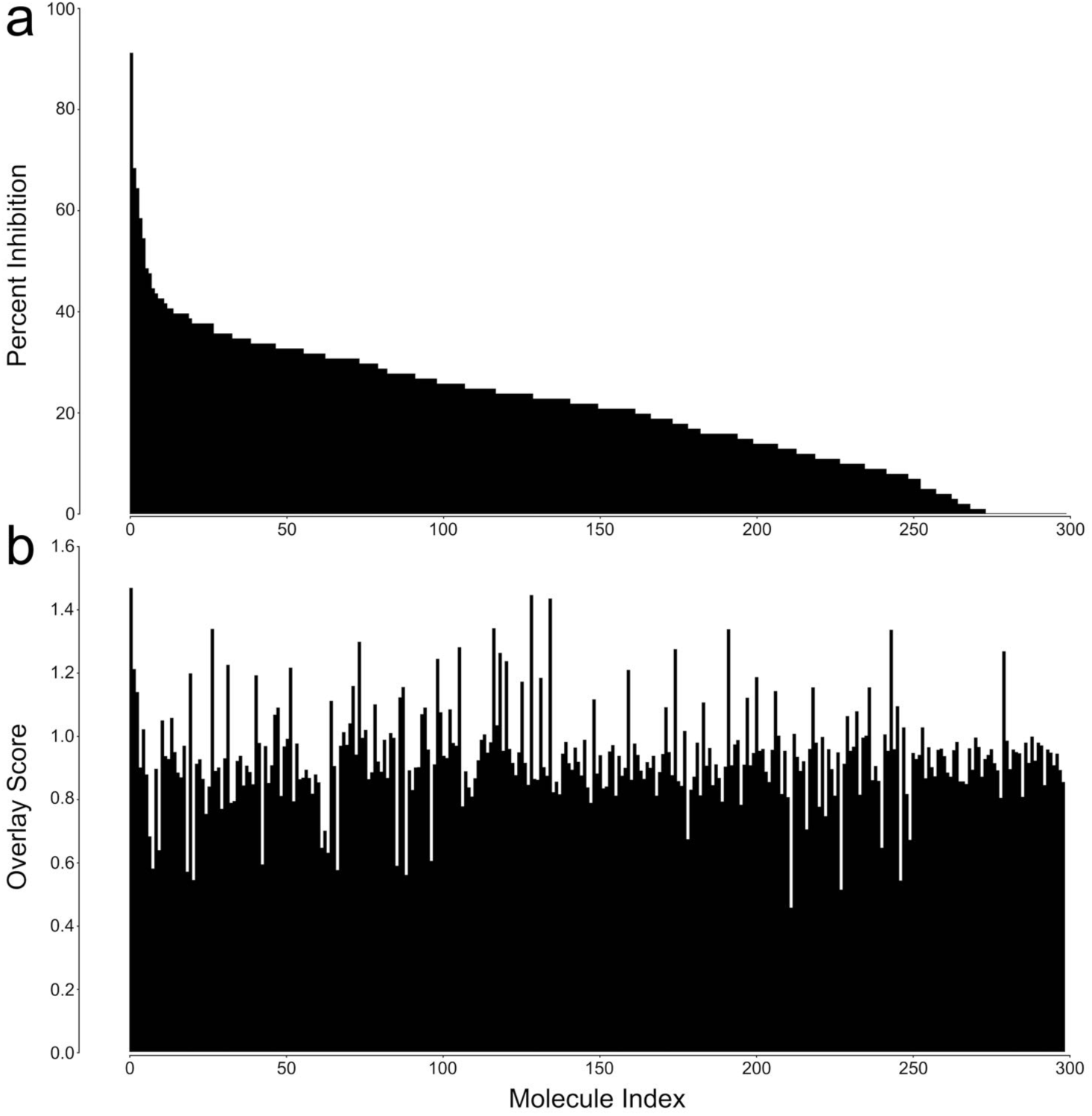
Quantitative comparison of EOG percent inhibition values for 3kPZS inhibitor candidates with their molecular similarity scores upon 3D overlay with 3kPZS. (a) The 299 assayed compounds were sorted by EOG activity values, from highest percent inhibition of 3kPZS response (left end of x-axis) to lowest (right end). (b) For these compounds shown in the same x-axis order as in (a), the ROCS TanimotoCombo molecular similarity scores following 3D flexible overlay with 3kPZS were plotted, equally weighting the electrostatic and volumetric components, with a maximum possible sum of 2.0. If the overlay similarity scores alone were highly predictive of activity, we would expect to see a pattern of high overlay scores corresponding to high percent inhibition values (that is, a similar profile of high scores decreasing to low scores, left to right, in (b) as well as (a)). However, the pattern of overlay scores in (b) is variable across the compounds, even for those with the highest percent inhibition values (TanimotoCombo and percent inhibition are virtually uncorrelated, with a Pearson correlation coefficient of 0.07). While for most hypotheses, only compounds with reasonably high overlay scores were assayed (meaning we pre-selected for overall molecular similarity), the data in (b) shows that overlay scores alone are not enough to predict the ability of a compound to inhibit 3kPZS activity. This drove the development of the tools in Screenlamp for identifying functional group patterns associated with biological activity (hypothesis-driven screening).

These results are consistent with a high degree of molecular similarity from overlay with a known ligand (or complementarity with the protein) being a useful feature in molecules that compete with a known ligand for receptor binding. However, overall similarity with a known ligand is typically insufficient to ensure that the same biological response is generated (e.g., activation or inhibition). Bioactive molecules form an exquisitely selective set of interactions in order to exclude the possibility of potentially lethal binding by the wrong ligands. The key is to identify which groups are making those specificity-determining interactions. Based on the experimental EOG data for 299 tested compounds, it is apparent that using hypothesis-driven functional group matching criteria in addition to ROCS-based similarity scoring yields greater enrichment of activity (Fig. 9), while also allowing identification of those critical groups. For instance, while retrieving compounds matching 3kPZS with a ROCS TanimotoCombo similarity score of 1.03 or more recovered 4 of the 5 most active molecules (with at least 50% inhibition), this set of retrieved molecules also included many (161) non-active molecules (Fig. 9b). Including additional selection criteria, such as the presence of a steroidal substructure, a sulfur or phosphorus overlay with the 24-sulfur atom in 3kPZS, and three sulfate oxygen matches, also yields 4 active molecules but only 13 inactives (Fig. 9a). Of the set of 299 molecules that were assayed, one molecule (PZS, or ZINC72400307, the most active), ranked highly (19^th^) when the TanimotoCombo score alone was considered. Hence, the selection of molecules for experimental assays would have been different if ROCS were used without additional hypothesis-based filtering criteria. However, this is expected, as the top 100 molecules ranked by different scoring functions (even those with similar features) in large database screening are often non-overlapping, due to the close similarity (virtual continuity) in score values, given a large number of molecules [2]. These results support that hypothesis-based chemical group filtering criteria, as facilitated by Screenlamp, not only decrease computational costs by appropriately reducing the chemical search space, but also are valuable for increasing the rate of retrieval of active compounds.

**Fig. 9.**
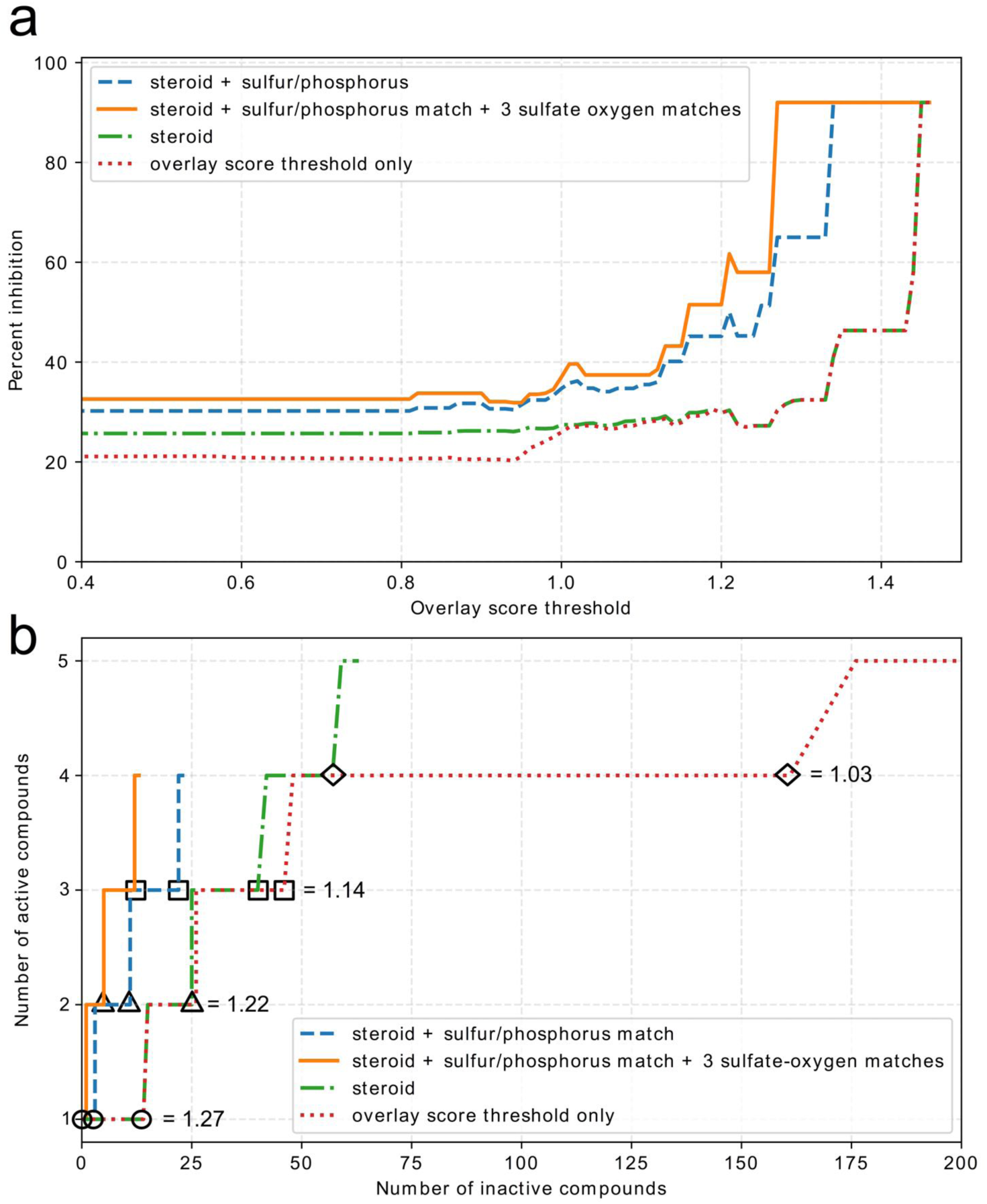
(a) Enrichment graphs showing the percent inhibition as a function of chemical and volumetric similarity of candidate molecules to 3kPZS, based on inhibition assays for 299 molecules selected by structure-activity hypotheses. The overlay score threshold refers to the ROCS TanimotoCombo score, equally weighting volumetric and pharmacophore similarity. The different traces on the graphs compare the enrichment for several hypotheses relative to using 3D similarity (overlay score) alone. (b) Receiver operating characteristic curve (rate of retrieval of true positives vs. false positives), with triangles, square and circle symbols used to show the point on each curve corresponding to a given score threshold, for cases in which the overlay score alone was used to select candidate compounds, versus when the overlay score was augmented by increasingly selective steroid-based hypotheses, resulting in fewer molecules being tested but a greater enrichment in active compounds. The curves show the number of active molecules (defined as having at least 50 percent inhibition of 3kPZS in experimental assays) for different overlay thresholds.

## Conclusions

Incorporating a hypothesis-driven strategy in computational and organismal biology has identified an inhibitor which in low concentration virtually nullifies the olfactory response of sea lamprey to the major mating pheromone, 3kPZS. Other highly active compounds were identified and are showing great promise as mating behavioral deterrents in ongoing stream trials. The ligand-based screening with Screenlamp also provided a series of simpler, nonsteroidal compounds that are significantly active and provide useful structure-activity information. To our knowledge, this presents the first successful application of structure-based drug discovery techniques to identify potent lead compounds for aquatic invasive species control. To enable other projects to benefit from this scalable, hypothesis-driven strategy which works easily with very large datasets, we have documented and are distributing the Screenlamp toolkit free of charge. (See Methods section on *Development of Screenlamp* for details.)

## Acknowledgements

We thank Qinghua Yuan for her contributions to the homology modeling of SLOR1 and Stacey Kneeshaw for evaluating protein-ligand energy minimization protocols for SLOR1-3kPZS docking and analyzing charge distributions for matching functional groups. This research was supported by funding from the Great Lakes Fishery Commission from 2012-present (Project ID: 2015_KUH_54031). We gratefully acknowledge OpenEye Scientific Software (Santa Fe, NM) for providing academic licenses for the use of their ROCS, OMEGA, QUACPAC (molcharge), and OEChem toolkit software.

## Supplementary Material

The CAS IDs of the 84 commercially available 3kPZS analogs (Table S1) and 2995 steroid substructure-containing compounds identified via SciFinder (Table S2) are provided in the Supplementary Material files 1 (supplementary_material_1_3kpzs_analogs.xlsx) and 2 (supplementary_material_2_cas_steroids.xlsx), respectively.

The TanimotoCombo similarity score distribution from overlaying the low-energy conformers of 3kPZS with the single-conformer drug-like database from ZINC (Fig. S1) and the molecular structure of PAMS-24 (Fig. S2) are available in Supplementary Material file 3 supplementary_material_3_suppl_figures.pdf). Structural validation reports on the SLOR1 model are provided as Supplementary Materials 4, 5, and 6 (supplementary_material_4_procheck_validation_slor1.pdf, supplementary_material_5_swissmodel_report_slor1_from_2vt4.pdf, and supplementary_material_6_swissmodel_2vt4_as_slor1_templ.pdf). The homology model of SLOR1 (file supplementary_material_7_slor1.pdb) and the binding mode of 3kPZS predicted in the orthosteric site of SLOR1 by flexible docking (file supplementary_material_8_3kpzs_in_slor1_orient.pdb) are available as individual PDB-format files and a PyMOL graphics file (supplementary_material_9_slor1_3kpzs_model.pse) in Supplementary Materials 7, 8, and 9, respectively. Also available are the PDB formatted structure of the CholMine cholate orientation (Supplementary Material 10, supplementary_material_10_cholate_orient_by_cholmine.pdb) compared with the SLIDE docking of 3kPZS (Supplementary Material 11, supplementary_material_11_3kpzs_docked_to_slor1_cholmine.pdb) in complex with the structure of SLOR1 in the CholMine reference frame (Supplementary Material 12, supplementary_material_12_slor1_3kpzs_cholmine_ref_frame.pdb), and a comparison of the predicted cholate and 3kPZS binding modes in SLOR1 as a PyMOL graphics file (Supplementary Material 13, supplementary_material_13_cholmine_cholate_pred_rel_to_3kpzs.pse). Three-dimensional overlays of the 15 most active antagonists of 3kPZS overlayed on the best-matching conformer of 3kPZS are provided as a PyMOL graphics file in Supplementary Material file 14 (supplementary_material_14_top15_cmpd_overlays.pse). A table containing experimental assay data for the 299 molecules tested (Table S3) is available in Supplementary Material file 15 (supplementary_material_14_percent_inhibition_299_candidates.xlsx).

